# Suppressor of Cytokine Signaling 3 Derived Peptide as a Therapeutic for Inflammatory, and Oxidative Stress Induced Damage to the Retina

**DOI:** 10.1101/2023.09.04.556227

**Authors:** Chulbul M. Ahmed, Anil P. Patel, Howard M. Johnson, Cristhian J. Ildefonso, Alfred S. Lewin

## Abstract

**Purpose:** Inflammation and oxidative stress are contributing factors to age-related macular degeneration (AMD) and other retinal diseases. We tested a cell penetrating peptide from the kinase inhibitory region of intracellular checkpoint inhibitor Suppressor of Cytokine Signaling 3 (R9-SOCS3-KIR) peptide for its ability to blunt the inflammatory or oxidative pathways leading to AMD.

**Methods:** We used Anaphylatoxin C5a to mimic the effect of activated complement, lipopolysaccharide (LPS) and TNFα to stimulate inflammation, and paraquat to induce mitochondrial oxidative stress. We used a human RPE cell line (ARPE-19) as proliferating cells and a mouse macrophage cell line (J774A.1) to follow cell propagation by microscopy or cell titer assays. We evaluated inflammatory pathways by monitoring nuclear translocation of NF-κB p65 and MAP kinase p38, and we used qRT-PCR and Western blots to evaluate induction of inflammatory markers. In differentiated ARPE-19 monolayers, we evaluated the integrity of tight junction proteins by microscopy and measurement of transepithelial electrical resistance. We used intraperitoneal injection of sodium iodate to test the ability of R9-SOC3-KIR to prevent RPE and retinal injury as assessed by fundoscopy, Optical Coherence Tomography (OCT) and histology.

**Results:** R9-SOCS3-KIR treatment suppressed C5a-induced nuclear translocation of the NF-kB activation domain p65 in undifferentiated ARPE-19 cells. TNF-mediated damage to tight junction proteins in RPE and the loss of transepithelial electrical resistance were prevented in the presence of R9-SOCS3-KIR. R9-SOCS3-KIR prevented the increased expression of genes related to inflammation in response to C5a treatment. R9-SOCS3-KIR also blocked lipopolysaccharide (LPS) induction of cyclooxygenase and inflammatory markers including IL-6, MCP1, COX-1 and IL-1β. R9-SOCS3-KIR prevented paraquat mediated cell death and enhanced the levels of antioxidant effectors. Daily eye drop instillation of R9-SOCS3-KIR protected against retinal injury caused by i.p. administration of sodium iodate.

**Conclusion:** R9-SOCS3-KIR blocks the induction of inflammatory signaling in cell culture and reduces retinal damage in a widely used model of RPE/retina oxidative injury. Since this peptide can be administered by corneal instillation, this treatment may offer a convenient way to slow the progression of ocular diseases arising from inflammation and chronic oxidative stress.

## Introduction

Age-related macular degeneration (AMD) is a leading cause of loss of central vision affecting nearly 50 million people worldwide, and, with the aging population, its incidence will increase in the coming years (1). Inflammation and oxidative stress are major drivers in the onset and propagation of AMD (2). Accumulation of oxidized lipoproteins as well as free radicals in retinal and choroidal tissue leading to chronic oxidative injury result in a full-blown inflammatory process, which, in turn, enhances the oxidative injury, thus self-perpetuating this vicious cycle. An important characteristic of AMD is the presence of drusen, extracellular proteolipid deposits located between RPE and Bruch’s membrane (3). Drusen contain byproducts of local inflammation and complement activation in addition to oxidized lipid and carbohydrate waste products (4). Continual injury to the RPE layer results in a compromised blood-retinal barrier that allows the cytokines and chemokines to enter and activate choroidal dendritic cells.

Advanced AMD presents as a ‘dry’ form of the disease, known as geographic atrophy, and leads to progressive and irreversible loss of central vision. The ‘wet’ form of the disease, characterized by choroidal neovascularization (CNV), occurs in about 10% of AMD patients, and may cause a sudden loss of central vision (4–7). CNV may respond to anti-vascular endothelial growth factor (VEGF) treatments delivered by intravitreal injection (8). Blocking VEGF pathway has its own drawbacks (9), and it does not prevent the ongoing inflammatory damage.

An overactive complement system is a critical mediator of AMD, diabetic retinopathy and Stargardt disease (4, 10, 11). Anaphylatoxins C5a and C3a, produced by the alternative complement pathway are detected in the drusen as well as in the serum of AMD patients (10, 12–14). The Y402H variant of complement factor H (CFH), a serum protein that normally tempers the alternative complement pathway, increase the progression of AMD (15, 16). The United States Food and Drug Administration, FDA, has approved two complement inhibitors for treatment of AMD, pegcetacoplan, a complement C3 inhibitor and avacincaptad pegol, a C5 inhibitor. Both treatments require repeated intravitreal injections.

A set of intracellular checkpoint protein inhibitors known as <u>s</u>uppressor <u>o</u>f <u>c</u>ytokine <u>s</u>ignaling (SOCS) suppress the expression of cytokines (17–19). Amongst these, SOCS1 and SOCS3 control JAK/STAT and toll like receptor (TLR) signaling, and represent important regulators of the extent and duration of an immune response. SOCS1 and SOCS3 are present in most somatic cells and allow cross talk between somatic cells and immune cells, suggesting an interconnected nature of their regulatory roles. Despite their importance in modulating inflammation, SOCS proteins have received less attention than the program death 1 cell protein (PD-1) and its ligand PD-1L, and cytokine T lymphocyte antigen 4 (CTLA-4) and its receptor (20). Evidence suggests that SOCS1 signaling is mediated by STAT1, while SOCS3 acts through STAT3 (18, 19, 21–23). A majority of cytokines and growth factors leading up to AMD act through STAT3 (24, 25); hence, we reasoned that use of a SOCS3 mimetic may be beneficial in suppressing inflammation leading to AMD. For SOCS1, we have shown that the kinase inhibitory region of the protein (KIR) by itself is sufficient to achieve the inhibition of the target kinases (26, 27). The native protein contains a SOCS box that is responsible for the proteasomal degradation of the SOCS1. Since, the KIR region of the peptide lacks the SOCS box, it is likely to sustain more cycles of inhibition of its target. In fact, it was shown for the SOCS3 protein that while the endogenous protein had a half-life of 0.7 hr, a cell penetrating peptide of the SOCS3 without the SOCS box had a half-life of 29 hr (28). Herein, we present data with a cell penetrating peptide containing the kinase inhibitory region (KIR) of SOCS3 (abbreviated as R9-SOCS3-KIR). We established the protective role of R9-SOCS3-KIR in response to activated complement, LPS and TNFα and oxidative damage in cell culture models. In a mouse model of acute oxidative stress induced by injection of sodium iodate, eye drop instillation of R9-SOCS3-KIR prevented retinal injury.

## Materials and Methods

### Cell Culture

ARPE-19 cells (CRL-2302) were obtained from ATCC (Manassas, VA) and grown in 1:1 mixture of Dulbecco’s Modified Eagle Medium and Hams F12 nutrient mixture (DMEM/F12, Sigma-Aldrich, St. Louis, MO) plus 10% fetal bovine serum (FBS), 1% each of penicillin and streptomycin in a humidified incubator at 37 °C and 5% CO_2_. Where indicated, cells were grown in 1% FBS (low serum) or serum free media. Murine macrophage cell line, J774A.1 (ATCC, TIB-67) was grown in Dulbecco’s Modified Eagle’s Medium (DMEM) with 10% fetal bovine serum (FBS) and 1% each of penicillin and streptomycin. Identity was certified by ATCC, and the cells were frozen immediately after their first passage, and passaged no more than three times for use in these experiments.

### Peptide Synthesis

The following peptides were chemically synthesized to 95% purity by GenScript (Piscataway, NJ). The peptide R9-SOCS3-KIR, with the sequence RRRRRRRRRLRLKTFSSKSEYQLVV consisted of the residues 20-35 from the kinase inhibitory region (KIR) of SOCS-3. The nine arginine residues on the N-terminus were included to allow penetration across biologic membranes. An inactive control peptide with the scrambled version of the KIR sequence with the following sequence RRRRRRRRRKSQYVRLSVLFEKTSL was used as a control. The peptides were dissolved in phosphate buffered saline (PBS) for use.

### NF-kB Promoter Assay

The plasmid, pNF-kB-Luc that has the NF-kB promoter linked to firefly luciferase and a plasmid that constitutively expresses Renilla luciferase from the thymidine kinase promoter (pRL-TK-Luc) were purchased from Promega (Madison, WI). pRL-TK-Luc serves as an internal control to test the efficiency of transfection. ARPE-19 cells were seeded at 80% confluency in 12 well plates and grown overnight. Cells, in triplicate, were placed in serum-free medium and treated with R9-SOCS3-KIR or the control peptide (both at 20 μM) for 1 hr, followed by treatment with C5a peptide (AbCam) at 50 ng/ml for 4 hrs. Cells were transfected using 2 μg of pNF-kB-Luc and 10 ng of pRL-TK-Luc per well using Lipofectamine (Invitrogen, San Diego, CA) for 24 hrs. The dual luciferase assay kit was obtained from Promega, and the firefly and Renilla luciferase activities were measured in cell lysates using a luminometer, following the manufacturer’s instructions.

### Immunohistochemistry

ARPE-19 or J774A.1 cells were seeded at 80% confluency in eight well chamber slides and grown overnight. They were placed in serum-free medium and treated with indicated concentrations of the R9-SOCS3-KIR or the control peptide for 1 hr, followed by 50 ng/ml of C5a peptide (Abcam) for 0.5 hr. Cells were washed and fixed with 4% paraformaldehyde for 30 min at room temperature and washed with PBS, followed by permeabilization with 1% Triton X-100 in PBS for 30 min at room temperature. Cells were then blocked in 10% normal goat serum in PBS containing 0.5% Triton X-100 followed by washing in 0.2% Triton X-100 in PBS (wash buffer). Rabbit polyclonal antibody for the NF-kB p65 subunit (Cell Signaling Technology) at 1:200 dilution was added and incubated overnight at 4 °C, followed by washing four times with the wash buffer. Cy-3 conjugated anti-rabbit secondary antibody (Invitrogen, 1:300 dilution) was added and incubated for 0.5 hr, followed by washing four times. Slides were co-stained with 4′,6’-diamidino-2-phenylindole (DAPI). After addition of the mounting media, cells were cover slipped and imaged in Keyence BZ-X700 fluorescence microscope. In other experiments, *E. coli* lipopolysaccharide (LPS), or paraquat (both from Sigma-Aldrich) were used at the concentrations indicated. Antibodies to phospho-STAT3 and phospho-p38 were obtained from Cell Signaling. Alexa-488 conjugated secondary antibody was from Thermo Fisher. For obtaining differentiated ARPE-19 cells, cells were grown in 1% FBS containing media with twice weekly change of media for 4 weeks until the cells had acquired a cuboidal morphology.

They were treated with R9-SOCS3-KIR or the control peptide at 20 μM for 3 hrs, followed by incubation with C5a 50 (ng/ml) or TNFα (10 ng/ml) for 48 hr. The processing of the cells after that was similar to that as described above. Cells were then incubated overnight with an antibody to ZO-1 (Invitrogen), followed by Cy-3 conjugated secondary antibody. The steps for processing further were similar to those described earlier (26).

### Measurement of Transepithelial Electrical Resistance (TEER)

ARPE-19 cells were seeded in 24-well transwell inserts (Griener Bio-one, 0.4 μm size) in 1% FBS containing media for ARPE-19 cells with twice weekly change of media for 4 weeks until the cells had started to look cuboidal. Cells were treated with 20 μM R9-SOCS3-KIR or the control peptide for 3 hrs followed by treatment with C5a peptide (50 ng/ml) or TNFα (10 ng/ml) for 48 hrs. Transepithelial electrical resistance (TEER) was measured using a voltohmmeter (EVOM2, World Precision Instruments), following the manufacturer’s instructions. The inserts were removed one at a time and placed in EVOM-2 chamber filled with the serum free media to take the TEER measurement. The value of a blank transwell filter was subtracted from each sample to get the net TEER.

### Enzyme-Linked Immunosorbent Assay (ELISA)

J774A.1 cells were seeded in 96 well plates at 80% confluency and grown overnight. They were placed in 1% FBS containing medium and treated with 20 μM R9-SOCS3-KIR or the control peptide for 1 hr followed by treatment with C5a peptide (50 ng/ml), followed by incubation overnight. Supernatants were harvested and used in triplicate to measure the concentration of VEGF-A by ELISA using the kit from PeproTech (Cranbury, NJ). In another experiment, following the peptide treatment of J774A.1 cells, LPS (1 μg/ml) was added, and cells were incubated overnight. Supernatants were harvested and used in triplicate to quantitate IL-1β using ELISA kit from PeproTech.

### Measurement of Released Nitric Oxide

J774A.1 cells were seeded at 80% confluence in 96 well plates and grown overnight. They were placed in 1% FBS containing medium and treated with increasing concentrations of R9-SOCS3-KIR or 30 μM of the control peptide for 1 hr, followed by addition of LPS (1μg/ml) and grown overnight. Supernatants were harvested and used in triplicate to measure the concentration of NO using Greiss reagent (Alexis Biochemicals, Plymouth Meeting, PA), following the manufacturer’s protocol. A standard curve was generated by using sodium nitrite as a substrate to determine the concentration in experimental samples.

### RNA extraction and qPCR

ARPE-19 or J774A.1 cells were seeded at 80% confluency in 12 well plates and grown overnight. ARPE-19 cells were transferred to serum free media and treated with 20 μM R9-SOCS3-KIR for 1 hr followed by treatment with C5a (50 ng/ml) for 4 hrs. J774A.1 cells were similarly placed in serum free media, treated with LPS (1 μg/ml) in one experiment and with paraquat (300 μg/ml) in another experiment for 4 hrs. Cells were washed with PBS, and total RNA was extracted using Trizol reagent (Invitrogen), and purified using the DirectZol RNA kit from Zymo Research (Irvine, CA). The i-Script kit from Bio-Rad (Hercules, CA) was used to generate cDNA. The quantitative polymerase chain reaction (qPCR) was carried out using a kit from Bio-Rad following the conditions described before (26). The sequence of primers used is provided in **Table 1**. The primers were synthesized by Eurofins (Louisville, KY). For the retinae obtained from mouse eyes, we employed a similar procedure. As an internal control, β-actin primers were used and the relative RNA concentrations were determined using the ^ΔΔ^Ct method (29).

**Table 1.**
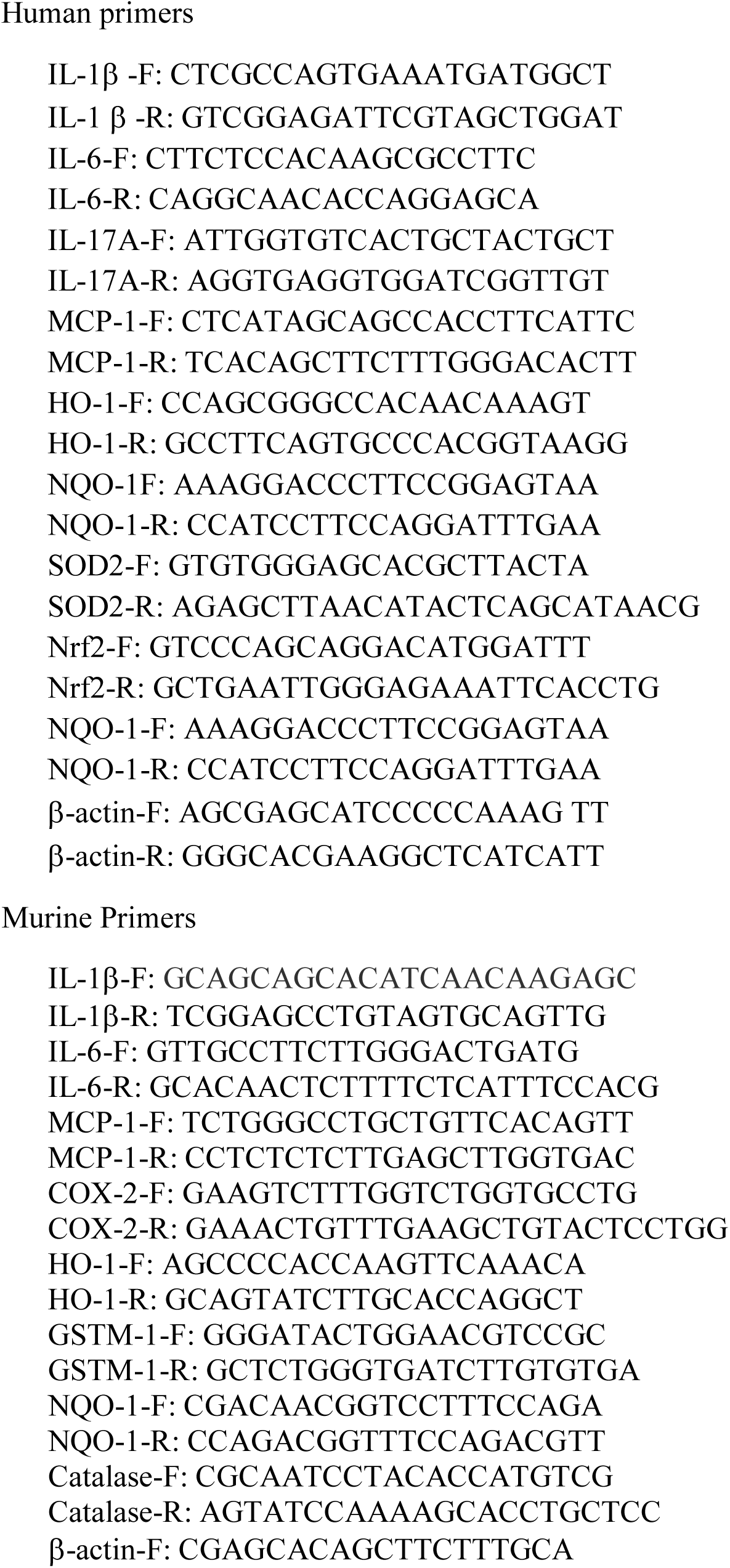
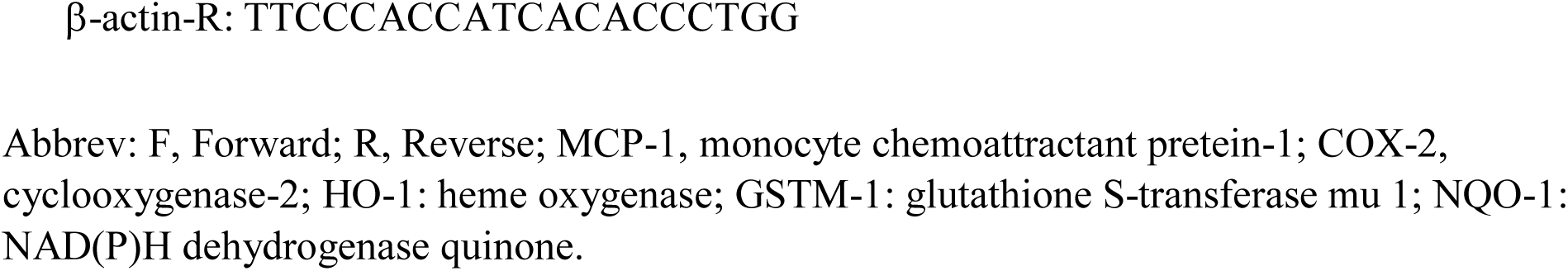
Nucleotide sequence of PCR primers used for qPCR.

### Western Blot Analysis

J774A.1 cells were seeded at 80% confluence in 6 well plates and grown overnight. They were transferred to 1% FBS containing medium and treated with R9-SOCS3-KIR or the control peptide for 1 hr, followed by treatment with LPS (1 μg/ml) and incubation overnight. Twenty micrograms of proteins from each sample were loaded on 12% polyacrylamide gel separated by electrophoresis, and then transferred to polyvinylidene difluoride (PVDF) membrane using an iBlot system (Thermo Fisher, Waltham, MA). The membrane was soaked in blocking buffer from LiCor Biosciences (Lincoln, NE) for 1 hr, followed by incubation with antibodies to COX-2 and α-tubulin (source of these reagents is provided in **Table 2**), as an internal control overnight. The membrane was rinsed four times with PBS containing 0.1% Tween 20. The appropriate IR dye-conjugated secondary antibodies (LiCor Biosciences; 1:5000 dilution in blocking buffer) was added for 0.5 hr. The membrane was washed four times with the blocking buffer and scanned with an Odyssey infrared imaging system (LiCor Biosystems). The experiment was repeated two more times using independent cell samples. Using the ImageJ system (NIH), the relative intensities of COX-2 and α-tubulin were calculated and plotted next to the first Western blot image. One-way ANOVA followed by Tukey’s test for multiple comparisons was used to determine the statistical difference in the relative intensities of bands under different treatments.

**Table 2.**
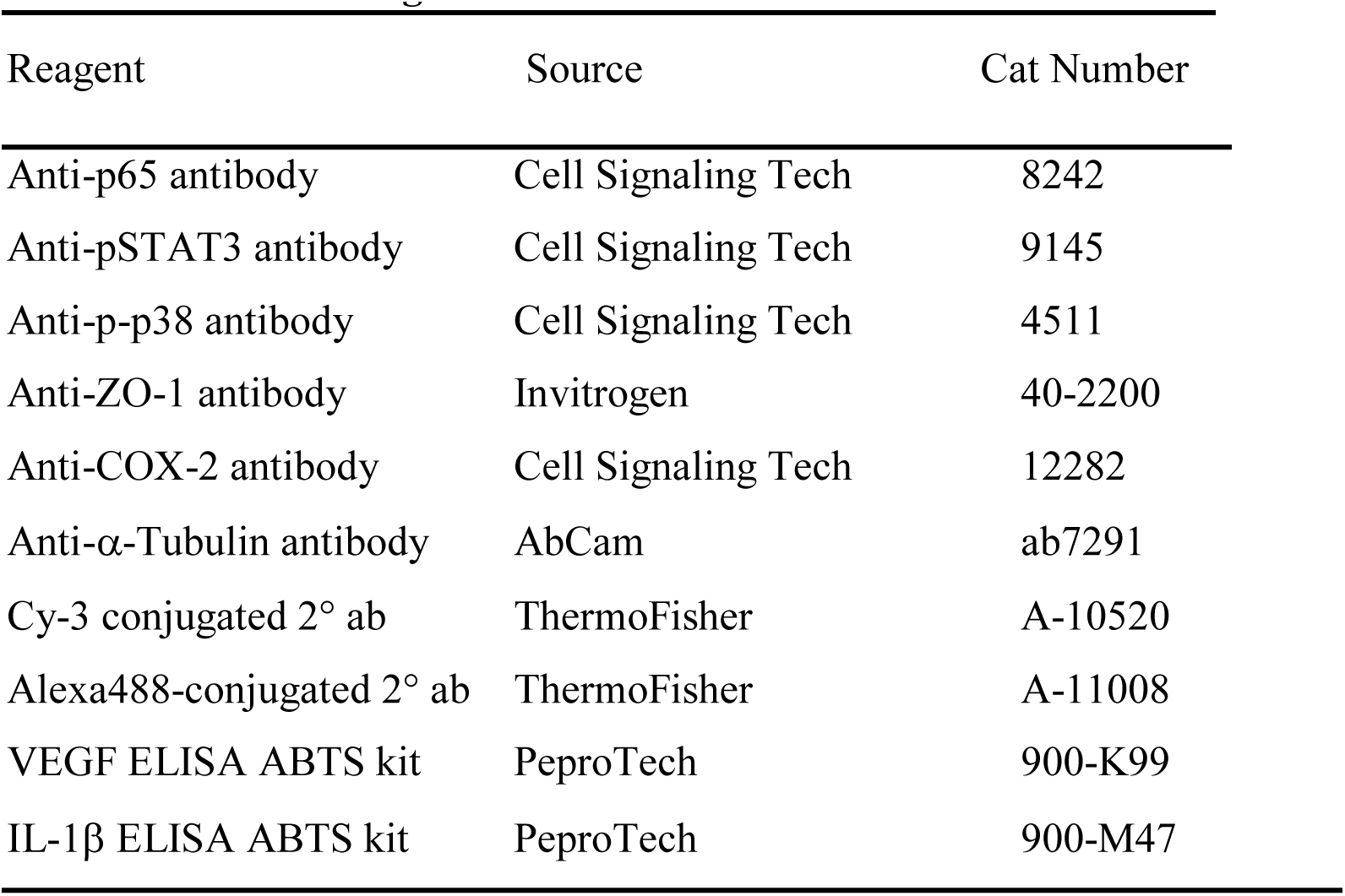
Source of reagents used.

| Reagent | Source | Cat Number |
| --- | --- | --- |
| Anti-p65 antibody | Cell Signaling Tech | 8242 |
| Anti-pSTAT3 antibody | Cell Signaling Tech | 9145 |
| Anti-p-p38 antibody | Cell Signaling Tech | 4511 |
| Anti-ZO-1 antibody | Invitrogen | 40-2200 |
| Anti-COX-2 antibody | Cell Signaling Tech | 12282 |
| Anti-α-Tubulin antibody | AbCam | ab7291 |
| Cy-3 conjugated 2° ab | ThermoFisher | A-10520 |
| Alexa488-conjugated 2° ab | ThermoFisher | A-11008 |
| VEGF ELISA ABTS kit | PeproTech | 900-K99 |
| IL-1β ELISA ABTS kit | PeproTech | 900-M47 |

### Sodium Iodate Induced Mouse Model of Oxidative Stress

The University of Florida Institutional Animal Care and Use Committee approved all procedures involving mice, and we conducted animal experiments in accordance with the Statement for the Use of Animals in Ophthalmic and Vision Research of the Association for Research in Vision and Ophthalmology (ARVO). C57BL/6 mice (BomTac, n=20, 10 male and 10 female) were divided into two groups, one for treatment with R9-SOCS3-KIR and the other with the control peptide. These mice had been tested to be sure that they did not carry the *Pde6β^rd1^* or the *Crb1^rd8^* mutations. The peptides were instilled in both eyes at 15 μg/eye in 2 μl once per day. Pre-treatment with these peptides started on day –1 and continued daily until day 3. On day 1, mice were injected i.p. with 25 mg/kg of sodium iodate (NaIO_3_), as described before (30). On day 3, fundoscopy and optical coherence tomography (OCT) were carried out. At the end of this, mice were humanely sacrificed and their retinae were harvested for RNA extraction and used for cDNA synthesis followed by qPCR as described above.

### Retinal Imaging and Cell Counting

Digital fundoscopy and spectral domain optical coherence tomography (SD-OCT) were carried, as described by us previously (27). To measure the thickness of outer nuclear layer (ONL), four measurements equidistant from the optical nerve were recorded from each eye. ONL thickness from both eyes was averaged and mean thickness for each group was calculated.

ImageJ software (https://imagej.nih.gov/ij) was used to count cells infiltrating in the vitreous. Area corresponding to the vitreous was marked and reflective spots corresponding to infiltrating cells were converted into binary images. To determine the number of cells in each area we used the count particles function of ImageJ, as described before (27). The number of infiltrating cells in both eyes of each mouse were averaged and the number of infiltrating cells in the R9-SOCS3-KIR or the control peptide groups were compared.

### Histopathology

One day 3 after NaIO_3_ administration, mice were humanely sacrificed. The eyes were dissected and placed in 4% paraformaldehyde overnight at 4 °C. Samples were washed in PBS, dehydrated using a graded series of alcohol solutions, and embedded in paraffin. Sectioning was carried out through the cornea-optic nerve axis at a thickness of 12 μm. Eight step sections were collected on different slides and stained with hematoxylin and eosin and the slides were prepared for microscopy.

### Statistical Analysis

Experiments from cell culture are presented as average ± standard deviation. We used a two-tailed Student’s *t*-test for unpaired data to compare the mean transcript levels. Results for the mice studies are presented as average ± standard deviation. For experiments that involved comparison of more than two samples, we used one-way analysis of variance (one-way ANOVA) followed by Tukey’s test to identify statistical significance, with GraphPad Prism 8.2.1 software (San Diego, CA). A p*-*value or adjusted p-value of less than 0.05 was considered statistically significant.

## Results

### R9-SOCS3-KIR dampens the response to activated complement

Activation of the alternate pathway of the complement system is a major contributor to the inflammatory response in AMD (4, 7). Given this critical role for activated complement, we tested the effect of anaphylatoxin, C5a on RPE-like cells to follow the effects on the pathways leading to inflammatory damage. After binding to its receptor, C5a acts through the NF-κB promoter (31, 32). Cross-talk of the C5a with TLR4 system also causes the activation of NF-κB pathway (33). To analyze the activation of NF-κB in RPE-like cells, we used a dual luciferase assay. Undifferentiated ARPE-19 cells were pre-treated with R9-SOCS3-KIR or its control peptide at 20 μM for 1 hr, followed by treatment with C5a peptide at 50 ng/ml for 4 hr. Treatment was followed by co-transfection with a firefly luciferase linked NF-kB promoter plasmid and a plasmid expressing Renilla luciferase under the control of thymidine kinase promoter, as an internal control. C5a and R9-SOCS3-KIR peptides remained in the culture for 24 hours before relative luciferase units were determined using the dual luciferase reporter kit from Promega (Madison, WI). C5a treatment caused a 9-fold induction of NF-κB promoter activity (**Figure 1A**). In the presence of R9-SOCS3-KIR, induction was reduced 75%. Treatment with C5a in the presence of the control peptide led to a 6.5-fold increase in activation of NF-kB promoter activity, indicating that the blockade of induction was dependent on the sequence of the peptide.

**Figure 1.**
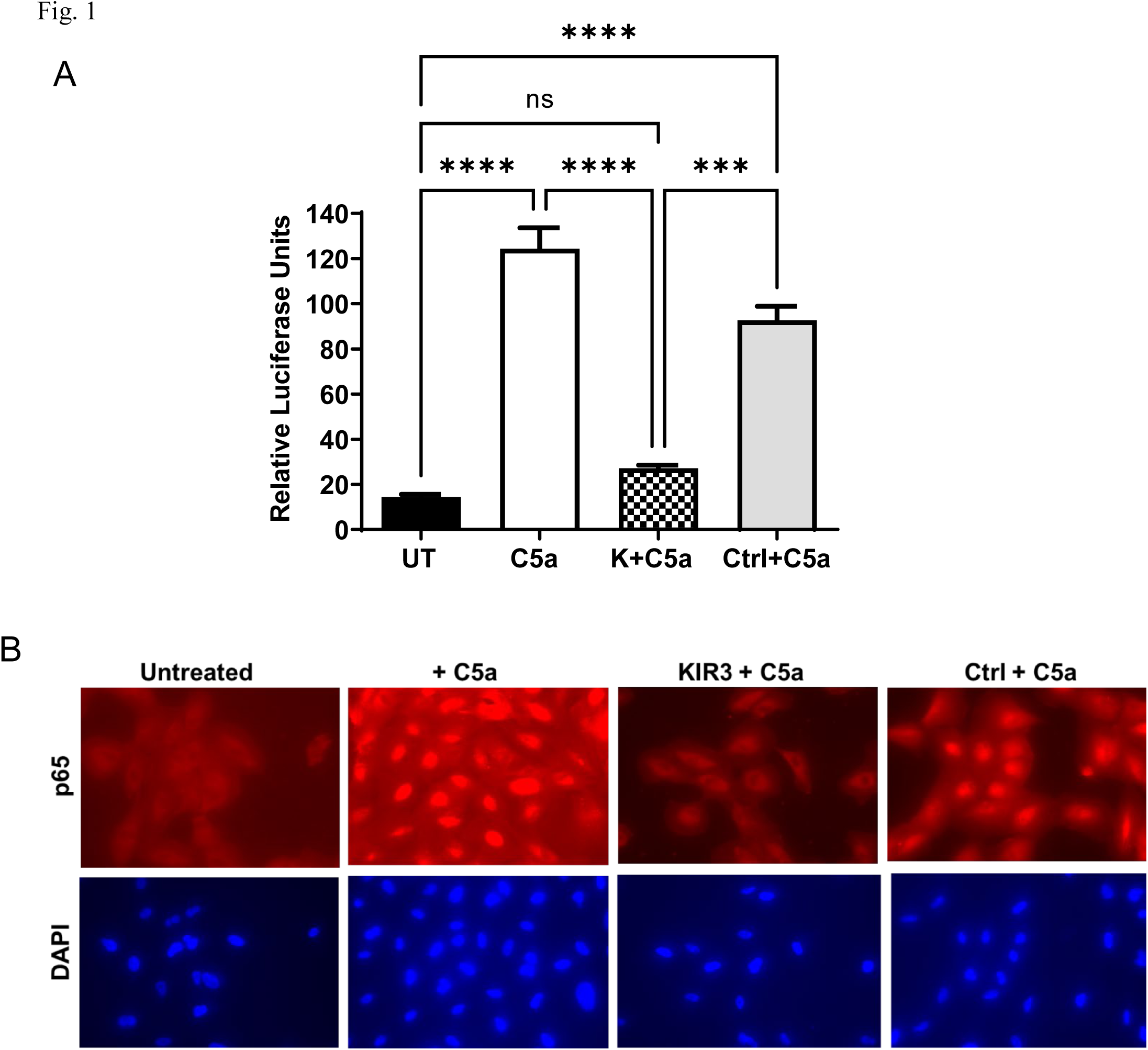
R9-SOCS3-KIR suppressed C5a mediated induction of NF-kB promoter. **A**. ARPE-19 cells in serum free medium were treated with R9-SOCS3-KIR or the control peptide (20 μM) for 1 hr followed by treatment with C5a (50 ng/ml) for 4 hr. They were then co-transfected with a mixture of plasmids with NF-kB promoter driven firefly luciferase and another plasmid with thymidine kinase driven Renilla luciferase as an internal control and grown for 24 hrs in 1% FBS containing medium. Cell lysates, were harvested and used to measure relative luciferase units using a Dual luciferase kit from Promega. **, p = 0.001. Error bars indicae the standard deviation. Abbrev: K, R9-SOCS3-KIR, Ctrl: R9-SOCS3-KIR-scrambled peptide. **B.** R9-SOCS3-KIR suppressed C5a-induced nuclear translocation of NF-κB subunit p65. ARPE-19 cells were grown overnight in 8 well plates, placed in serum-free medium and treated with R9-SOCS3-KIR or its control peptide at 20 μM for 1 hr, followed by addition of C5a (50 ng/ml) for 30 min. Cells were stained with an antibody to p65, followed by staining with Cy-3 conjugated secondary antibody and DAPI, and imaged in a fluorescence microscope using the same exposure time and light intensity. Scale bars equal 50 μm.

As an independent assay of NF-kB activation, we analyzed the nuclear translocation of the p65 subunit of NF-kB in response to complement treatment (**Figure 1B**). Treatment with C5a resulted in nuclear translocation of p65 in most cells, indicating the activation of NF-κB pathway. Cells that were pre-treated with R9-SOCS3-KIR followed by C5a, did not exhibit nuclear accumulation of p65, while the negative control peptide did not affect the translocation of NF-κB, indicating the specificity of action of R9-SOCS3-KIR.

Complement activation can lead to increased cell permeability and damage to the tight junctions between RPE cells. The loss of tight junction proteins results in decrease of transmembrane electrical resistance (TEER) by allowing the paracellular passage of ions from the basal to apical side of the cells (34). To determine whether R9-SOCS3-KIR could protect an RPE monolayer, ARPE-19 cells were grown in low serum media for 4 weeks until the cells had differentiated and acquired a cobblestone-like morphology. These cells were pre-treated with R9-SOCS3-KIR or the control peptide (20 μM) for 3 hr followed by treatment with C5a peptide (50 ng/ml) for 48 hrs. At the end of treatment with C5a, TEER was measured with a voltohmmeter (**Figure 2A**). C5a treatment caused a 72% decrease in TEER, as compared with untreated cells. In the presence of R9-SOCS3-KIR, the loss of TEER caused by C5a was only 16%, whereas the control peptide showed a 70% decrease in TEER, indicating a protective role of R9-SOCS3-KIR in maintaining the integrity of tight junction proteins in the presence of C5a.

**Figure 2.**
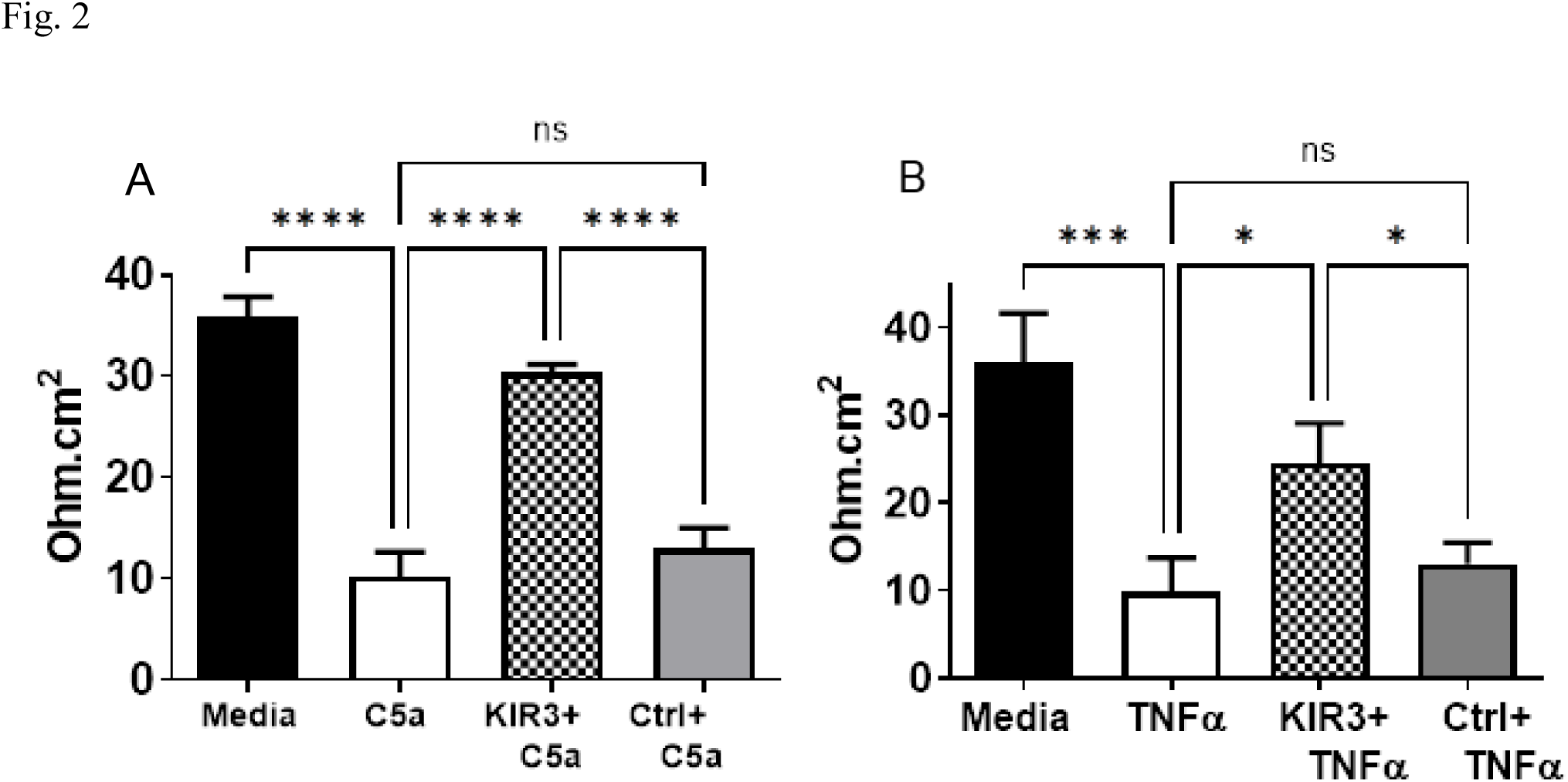
A. R9-SOCS3-KIR protected ARPE-19 cells against the loss of transepithelial electrical resistance (TEER) caused by C5a and TNFα. **A**. ARPE-19 cells grown in transwell plates (24 well) for four weeks in low serum until they attained a cobblestone morphology were treated with R9-SOCS3-KIR or control peptide (20 µM) for 2 hrs, followed by treatment with C5a (50 ng/ml) for 48 hr. TEER in individual wells, in triplicate, was measured using EVOM2 voltohmmeter. **B.** ARPE-19 cells were grown as described in **(A)** were treated with R9-SOCS3-KIR or its control peptide (both at 20 μM) for 3 hrs. TNFα at 10 ng/ml was added and cells were incubated for 48 hrs. TEER was measured in triplicate, with a voltohmmeter. Units are ohms/cm^2^. One-way analysis of variance (ANOVA), followed by Tukey’s multiple comparisons was used to test significance. *, p<0.05; ***, p<0.001, ****, p<0.0001. Abbreviations: KIR-3, R9-SOCS3-KIR, Ctrl, scrambled R9-SOCS-3 KIR peptide.

Increased levels of TNFα have been reported to contribute to choroidal neovascularization and to stimulate epithelial to mesenchymal transition in the RPE (35, 36). We therefore tested the effect of TNFα on RPE cells. Differentiated ARPE-19 cells that were grown in transwell plates were treated with TNFα, which decreased TEER by 70%. In the presence of R9-SOCS3-KIR, TEER decreased by only 20% (**Figure 2B**), while in the sample containing the control peptide TEER was not statistically different from TNFα alone, implying the protective role of R9-SOCS3-KIR in preserving the integrity of tight junction proteins.

To visualize the tight junctions among RPE cells, RPE monolayers, treated with C5a or TNFα as described above, were incubated with an antibody to the tight junction protein, zona occludens 1 (ZO-1), followed by staining with a Cy-3 conjugated secondary antibody and observed in a fluorescence microscope (**Supplemental Figure 1**). Untreated cells showed a continuous distribution of the ZO-1 protein along the neighboring cells indicating the integrity of the monolayer, as seen in the differentiated RPE cells. Treatment with either the C5a peptide or TNFα resulted in lack of uniform distribution of the tight junction protein, with marked disruptions between cells, as is expected of cells in stress (37). The presence of R9-SOCS3-KIR prevented the loss of ZO-1 along the cell membranes. In contrast, the control peptide did not prevent the loss of ZO-1 caused by C5a or TNFα.

To test the ability of R9-SOCS3-KIR to suppress the inflammatory markers induced by C5a, we measured transcript levels of inflammatory cytokines using qRT-PCR (**Figure 3A**). Treatment with C5a caused a 3 to 3.5-fold increase in the RNA levels of the cytokines IL-1β, IL-6, IL-17A and the chemokine MCP-1. However, these levels diminished to 1 to 1.5-fold when treated with R9-SOCS3-KIR, indicating the immunosuppressive effect of this peptide. Thus, R9-SOCS3-KIR is a potent inhibitor of C5a toxicity at multiple levels and is capable of dampening its effect.

**Figure 3.**
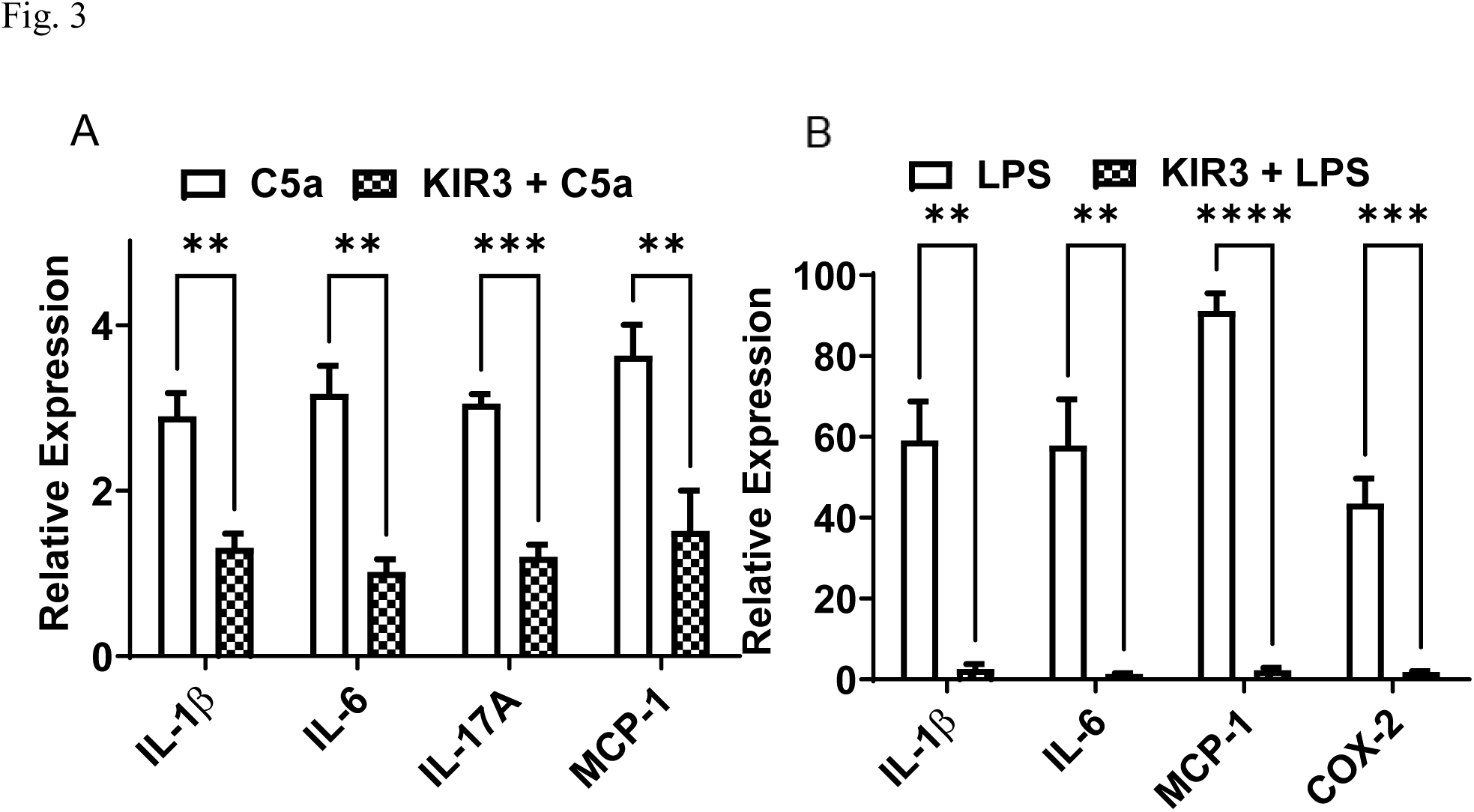
C5a and LPS induced inflammatory mediators were suppressed in the presence of R9-SOCS3-KIR. **A**. ARPE-19 cells were seeded in 6 well plates and grown overnight. They were placed in serum free media and treated with or without R9-SOCS3-KIR (20 μM) for 1 hr, followed by treatment with C5a peptide (50 ng/ml) for 4 hrs. Cells were harvested and RNA was extracted for cDNA synthesis. The cDNA was used for qPCR using the primers for the target genes indicated and β-actin primers as an internal control. The results represent the average of biological triplicates, and the error bar indicates the standard deviations. For each comparison, Student t test for statistical significance showed a p value of 0.001 between C5a and R9-SOCS3-KIR + C5a treatments. **B.** Anti-inflammatory effects of R9-SOCS3-KIR observed in LPS treated J774A.1 cells. Mouse macrophage cell line, J774A.1 cells were grown overnight in 12 well plates. They were placed in serum free media and treated with R9-SOCS3-KIR (20 μM) for 1 hr, followed by addition of LPS (1 μg/ml) for 4 hrs. Quantitative RT-PCR was carried out using primers for the target genes indicated and β-actin primers as internal control. The bars represent the average of triplicates ± sd. For each comparison, statistical significance was measured using Student’s t test. **, p <0.01; ***, p <0.001;****, p<0.0001.

### R9-SOCS3-KIR protects against the inflammatory damage caused by LPS

To evaluate the anti-inflammatory effects of R9-SOCS3-KIR, we used LPS, which activates NF-kB and MAP kinase p38 through the Toll-like receptor 4 (38, 39). We used a mouse macrophage cell line, J774A.1, as a surrogate for activated mononuclear phagocytes in the retina. Macrophages in particular are known to play a central role in progression to AMD (40), and diabetic retinopathy (41). To define the anti-inflammatory actions of R9-SOCS3-KIR in mononuclear phagocytes, we treated J774A.1 cells with R9-SOCS3-KIR or control peptide for 1 hr, followed by treatment with LPS (1 μg/ml) for 4 hr. RNA was extracted from these cells and converted into cDNA and used as a template for qPCR using the primers for the inflammatory genes indicated (**Figure 3B**). LPS treatment caused a 60-fold increase in IL-1β and 57-fold increase in IL-6 transcript levels, which was prevented in the presence of R9-SOCS3-KIR. Although the chemokine MCP-1 transcript was increased 90-fold following LPS treatment, it remained at baseline in the presence of R9-SOCS3-KIR. The transcript for the enzyme cyclooxygenase 2 (COX-2) went up 45-fold, which remained at the baseline level in the presence of R9-SOCS3-KIR. Thus, R9-SOCS3-KIR is a potent suppressor of pro-inflammatory gene expression.

To evaluate the impact of SOC3-KIR peptide on the nuclear translocation of NF-kB and p38 in macrophage cells, we pretreated J774A.1 cells in serum-free medium with R9-SOCS3-KIR or its control (both at 20 μM) followed by addition of LPS (1 μg/ml) for 30 min. Cells were washed and stained with primary antibodies to p65, the active subunit of NF-kB or the antibody to phospho-p38 (the activated MAP kinase p38), washed, stained with fluorescent secondary antibodies and DAPI to visualize nuclei (**Figure 4**). LPS treatment caused the nuclear translocation of NF-κB (**A**) and p-p38 (**B**). However, treatment with the R9-SOCS3-KIR peptide blocked the nuclear translocation of both p65 and p38, while the control peptide did not have a significant impact on either of them. These results suggest that the R9-SOCS3-KIR peptide blocks the induced nuclear translocation of NF-kB and p38 therefore explaining the strong suppression of proinflammatory gene expression.

**Figure 4.**
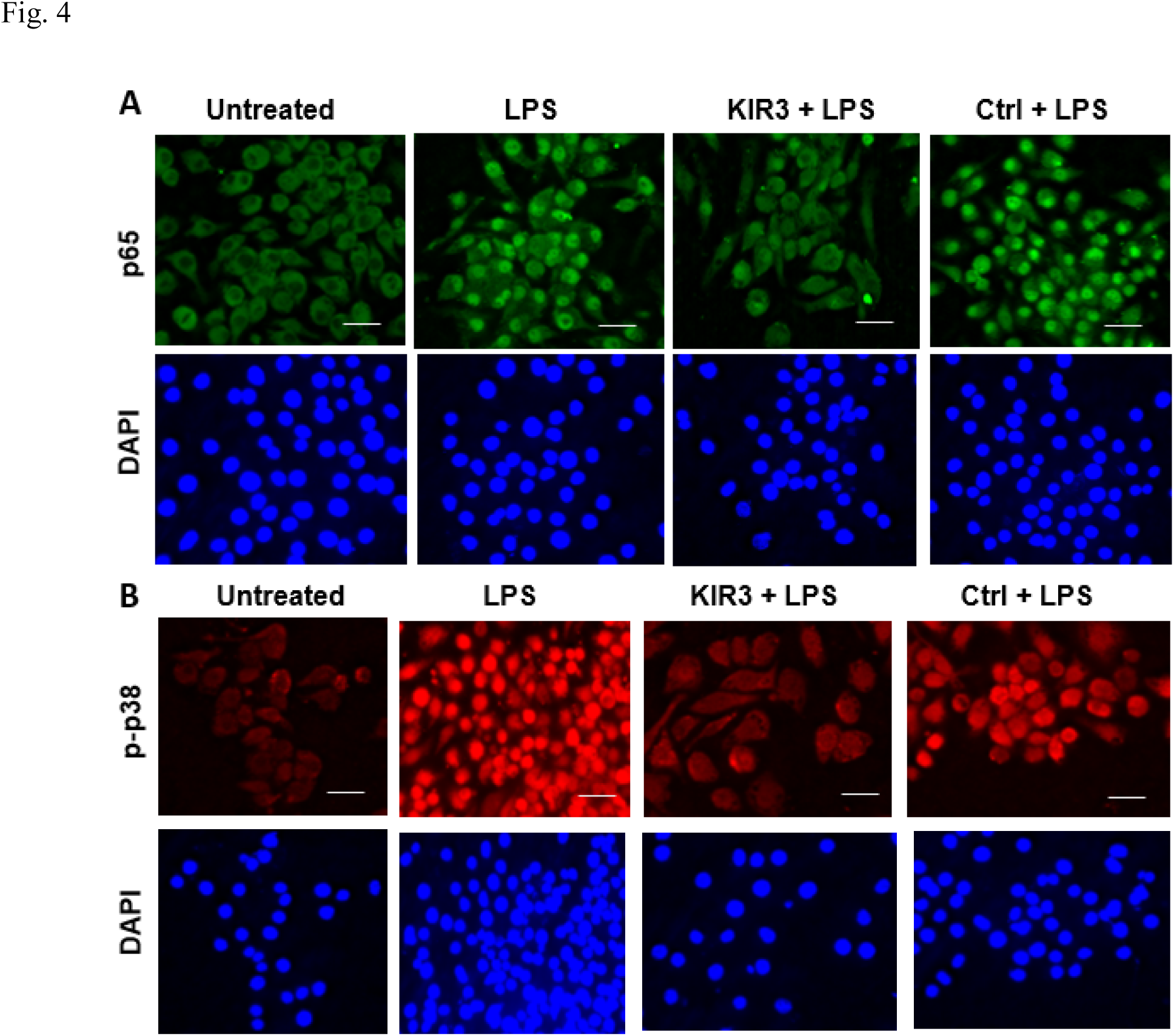
R9-SOCS3-KIR suppressed LPS induced nuclear translocation of NF-κB (A) and MAP kinase p38 (B) in J774A.1 cells. J774A.1 cells were seeded in 8-well cell culture slide and grown overnight. Cells were transferred to serum free medium and pre-treated with R9-SOCS3-KIR peptide, or its control peptide at 20 µM for 1 hr, followed by treatment with LPS (1μg/ml) for 30 min. Cells were stained with an antibody to p65, the active subunit of NF-κB (**A**), or p-p38 (**B**). Secondary antibodies conjugated to Alexa-488 (**A**, green), or Cy-3 red (**B**, red) and DAPI (nuclei, bottom row in A and B) were used and imaged in a fluorescence microscope. Scale bar, 50 μm.

Next, we characterized the impact of our peptide on mononuclear phagocyte inflammatory functions. To confirm the protective role of R9-SOCS3-KIR against the inflammatory effect of LPS, we pre-treated J774A.1 cells with 0, 3, 10 and 30 μg/ml of R9-SOCS3-KIR or 30 μM of the control peptide for 1 hr. This was followed by addition of LPS at 1 μg/ml and incubation overnight. The supernatants were harvested and used for the quantitation of nitric oxide (NO) using the Greiss reagent (**Supplemental Figure 2A**). There was a 16%, (p<0.01), 45% and 73% (p < 0.001) decrease in the release of NO in the presence of 3, 10 and 30 μg/ml of R9-SOCS3-KIR, respectively. In contrast, there was only a 4% decrease of NO in the presence of control peptide at 30 μg/ml, thus suggesting the dose-dependent inhibition of NO production in the presence of R9-SOCS3-KIR. We also studied the impact of our peptide in the secretion of proinflammatory cytokines using ELISA. J774A.1 cells that were pre-treated with 20 μg/ml of R9-SOCS3-KIR or its control peptide for 1 hr followed by treatment with LPS at 1 μg/ml, overnight were used to harvest the supernatants. Secretion of IL-1β in these supernatants was quantitated by ELISA (**Supplemental Figure 2B**). There was a negligible amount of IL-1β in the untreated cells. Treatment with LPS resulted in secretion of 1,600 pg/ml of IL-1β, which was reduced by 75% in the cells in the presence of R9-SOCS3-KIR, whereas the control peptide reduced this secretion by only 20% (p<0.0001 for both), which agrees with our qRT-PCR observations above (**Figure 3)**. We studied changes in the protein levels of another inflammatory marker, cyclooxygenase 2 (COX-2) by performing Western blots with cell lysates (**Supplemental 2 C**). J774A.1 cells were pre-treated with R9-SOCS3-KIR or its control peptide at 20 μM for 1 hr followed by treatment with LPS at 1 μg/ml for 24 hrs. Membranes were probed with antibodies to COX-2 and α-tubulin as an internal control. The relative intensities of COX-2 and the internal control α-tubulin were measured using image J (NIH) software. The relative intensities of COX-2 and α-tubulin (used as an internal loading control) in untreated cells was taken as 1, and the relative intensities of other treatments were plotted (**Supplemental Fig. 2D**). LPS treatment resulted in 6-fold increase in the synthesis of COX-2, which was reduced to 2.5-fold in the presence of R9-SOCS3-KIR, and was unaffected in the presence of control peptide, indicating that the R9-SOCS3-KIR peptide caused a reduction in expression of COX-2 protein. Overall, our studies support the hypothesis that R9-SOCS3-KIR mediated inhibition of STAT3 in mononuclear phagocytes can significantly inhibit their proinflammatory functions.

### R9-SOCS3-KIR reduced the secretion of VEGF-A by mononuclear phagocytes

Since invading macrophages exacerbate CNV by elevating VEGF, we tested the impact of R9-SOCS3-KIR on the mouse macrophage cell line. J774A.1 cells were pre-treated with R9-SOCS3-KIR or the control peptide for 1 hr followed by treatment with C5a (50 ng/ml) and incubation overnight. Cell supernatants were harvested and used for quantitation of VEGF-A by ELISA. Supernatants of untreated cells showed 18.4 ± 3.9 pg/ml, while treatment with C5a resulted in a 47-fold (p<0.001) increase in the VEGF-A level (**Supplemental Figure 3**). The presence of R9-SOCS3-KIR, led to an 80% decrease in level of VEGF-A (p < 0.001), while the control peptide treatment resulted in only a 20% decrease in VEGF-A levels (p < 0.001), showing that the reduction in VEGF-A secretion was specific to R9-SOCS3-KIR.

### R9-SOCS3-KIR alleviates the oxidative damage caused by paraquat

Since oxidative stress contributes to RPE damage and AMD pathogenesis (42), we measured the effect of SOCS3-KIR on RPE cells treated with paraquat to cause an increase in mitochondrial reactive oxygen species (43). We treated ARPE-19 cells with 20 μM of R9-SOCS3-KIR or the control peptide for 1 hr, followed by incubation with 300 μM of paraquat for 24 hrs. Cells were stained with hematoxylin and eosin and viewed in a confocal microscope (**Figure 5A**). Untreated cells showed a confluent monolayer. Treatment with paraquat resulted in disruption of this layer with gaping holes. Cells treated simultaneously with R9-SOCS3-KIR, were protected from this damage and remained confluent, while the control peptide did not prevent the damage to the cells. We assessed survival of ARPE19 cells by using the CellTiter Aqueous reagent (Promega) and reading the absorbance in a plate reader. Cell survival was calculated by using the absorbance in untreated cells as 100%. In the presence of 3, 10 and 30 μg/ml of R9-SOCS3-KIR, we noted a 20%, 50% and 80% survival of cells, respectively (p < 0.001 for all), suggesting a dose-dependent protection against damage caused by paraquat in these cells (**Figure 5B**). In the presence of the control peptide, only 7% of cells survived, suggesting the specificity of anti-oxidant effect with the R9-SOCS3-KIR peptide.

**Figure 5.**
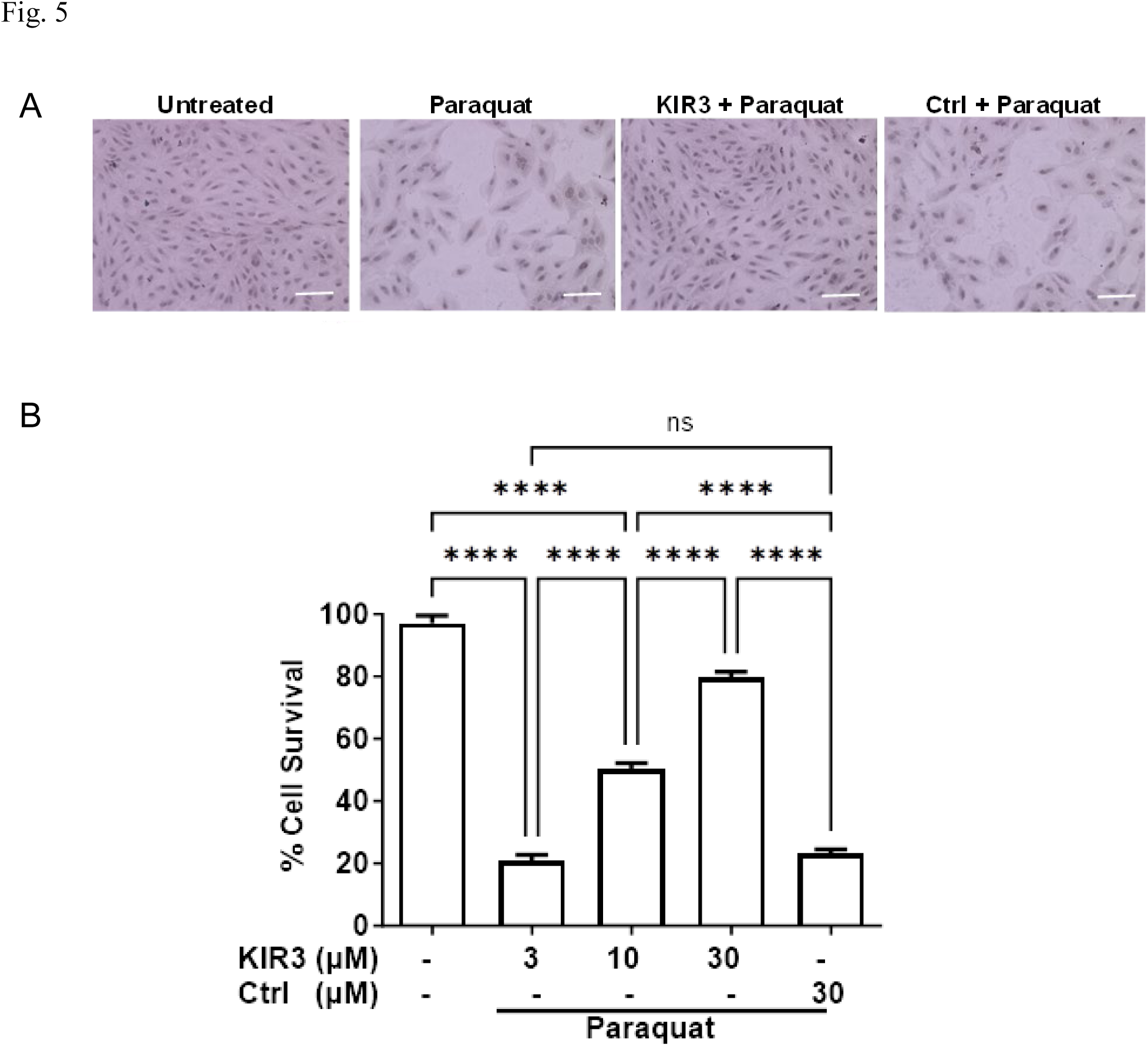
R9-SOCS3-KIR prevented paraquat mediated oxidative damage to RPE-like cells. **A**. ARPE-19 cells were seeded in 8 well slides and grown overnight before transfer to serum-free medium. They were pretreated with R9-SOCS3-KIR or its control peptide (both at 20 μM) for 1 hr followed by treatment with paraquat at 300 μM for 48 hrs. Cells were stained with hematoxylin and eosin and imaged with a confocal microscope. Scale bar: 50 μm. **B.** In cells grown and treated as in (**A**), cell survival was assessed by using CellTiter Aqueous reagent (Promega) and reading the absorbance in a plate reader. Absorbance in untreated cells was set as 100%, and survival in other treatments was calculated. One-way analysis of variance (ANOVA), followed by Tukey’s test for multiple comparisons. ****, p<0.0001.

To test if the protection against oxidative damage noted above was associated with an induction of anti-oxidant factors, we carried out qRT-PCR for effector molecules that have protective properties (**Figure 6**). We noted a 6.5-fold increase in heme oxygenase 1 (HO-1), which was induced 8.7-fold in the presence of R9-SOCS3-KIR. Similarly, the levels of SOD-2, Nrf-2 and NqO-1 were increased 3-fold by paraquat alone, and were induced further by 1.5-fold to 2-fold in the simultaneous presence of R9-SOCS3-KIR. The induction of these genes that are regulated by antioxidant response element (ARE), as well as the induction of Nrf-2, a transcription factor for antioxidant enzymes, is consistent with the survival of ARPE-19 cells seen above in the presence of paraquat.

**Figure 6.**
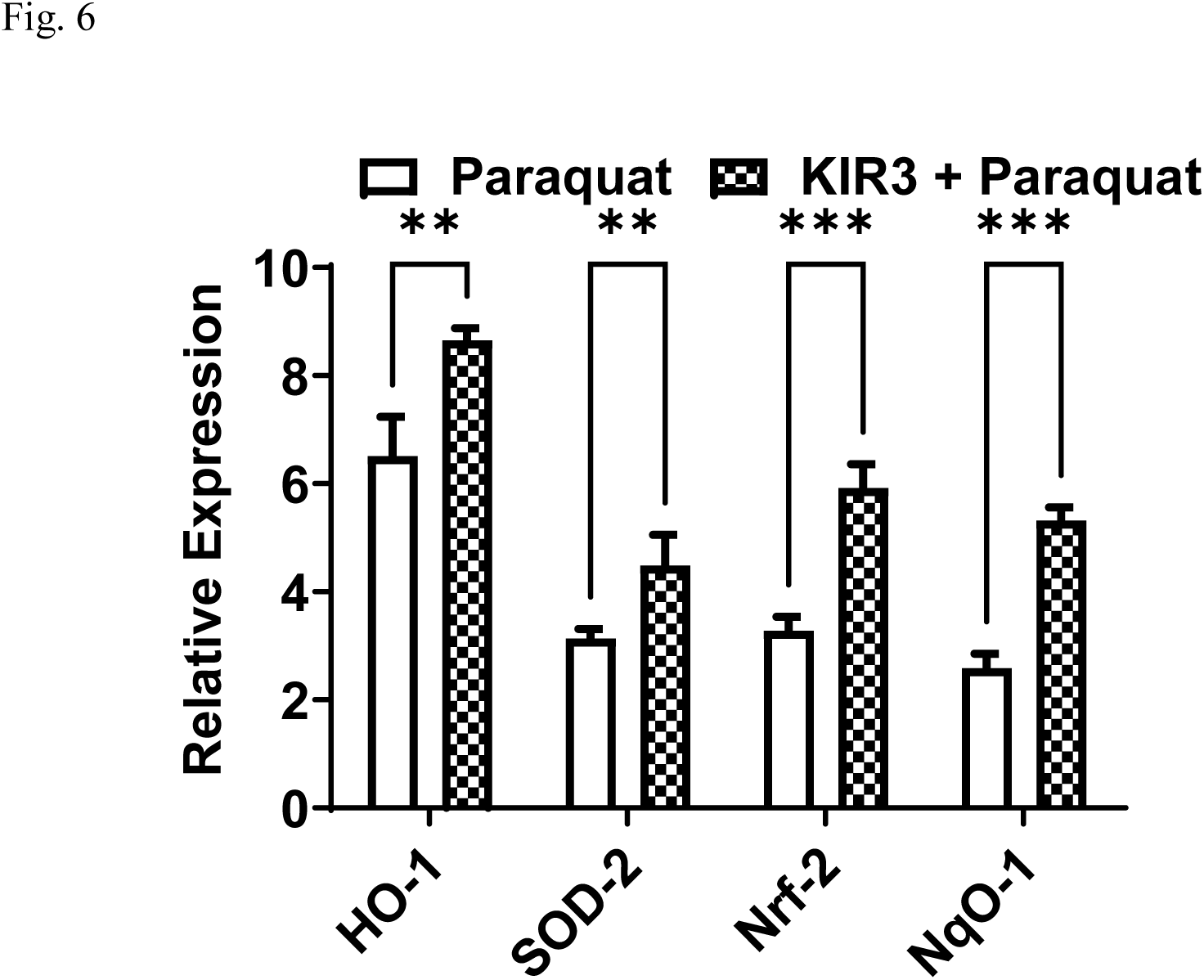
R9-SOCS3-KIR enhances antioxidant response in paraquat treated ARPE-19 cells. ARPE-19 cells were grown overnight in 12 well plates. They were placed in 1% FBS containing medium and treated with R9-SOCS3-KIR (20 μM) for 1 hr, followed by treatment with paraquat (300 μM) for 4 hr. Cells were washed and RNA was extracted and used for cDNA synthesis. Quantitative PCR was carried out using the primers for target genes indicated above and β-actin as an internal control. Units represent expression level relative to β-actin. Student’s t test for unpaired data was used to calculate the statistical significance in each set of samples. *, p <0.05; **, p <0.01, ***. p<0.001. Abbreviations: HO-1, heme oxidase 1; SOD2, superoxide dismutase 2, Nrf2, NFE2-like BZIP transcription factor 2, NQO-1, NAD(P)H Quinone Dehydrogenase 1.

### R9-SOCS3-KIR acts through STAT3 inhibition

There is evidence that SOCS1 inhibits signaling mediated by STAT1, while SOCS3 predominantly inhibits signaling through STAT3 (18, 44, 45). While SOCS1 binds directly to the kinase domain of specific target JAKs, TYKs or adaptors, SOCS3 docks on the gp130 subunit of IL-6 family of receptors and subsequently binds to the corresponding kinase domain of specific JAKs.

To test the effect of R9-SOCS3-KIR acting through STAT3, we treated ARPE-19 cells with IL-6. IL-6 and IL-17 are implicated in the progression of dry AMD to geographic atrophy (46) and CNV (47). Since IL-6 acts through STAT3 activation, we followed the effect of IL-6 on activated STAT3. ARPE-19 cells were treated with R9-SOCS3-KIR or the control peptide for 1 hr followed by addition of IL-6 (50 ng/ml) for 0.5 hr. Cells were incubated with an antibody to phospho-STAT3, the activated form of STAT3, followed by staining with Alexa-488 conjugated secondary antibody and DAPI. Fixed cells were imaged in a fluorescence microscope (**Figure 7**). Treatment with IL-6 resulted in nuclear translocation of pSTAT3 in ARPE-19 cells. This translocation was blocked in the presence of R9-SOCS3-KIR, but not by the vehicle control, thus showing the inhibition of STAT3 signaling by R9-SOCS3-KIR.

**Figure 7.**
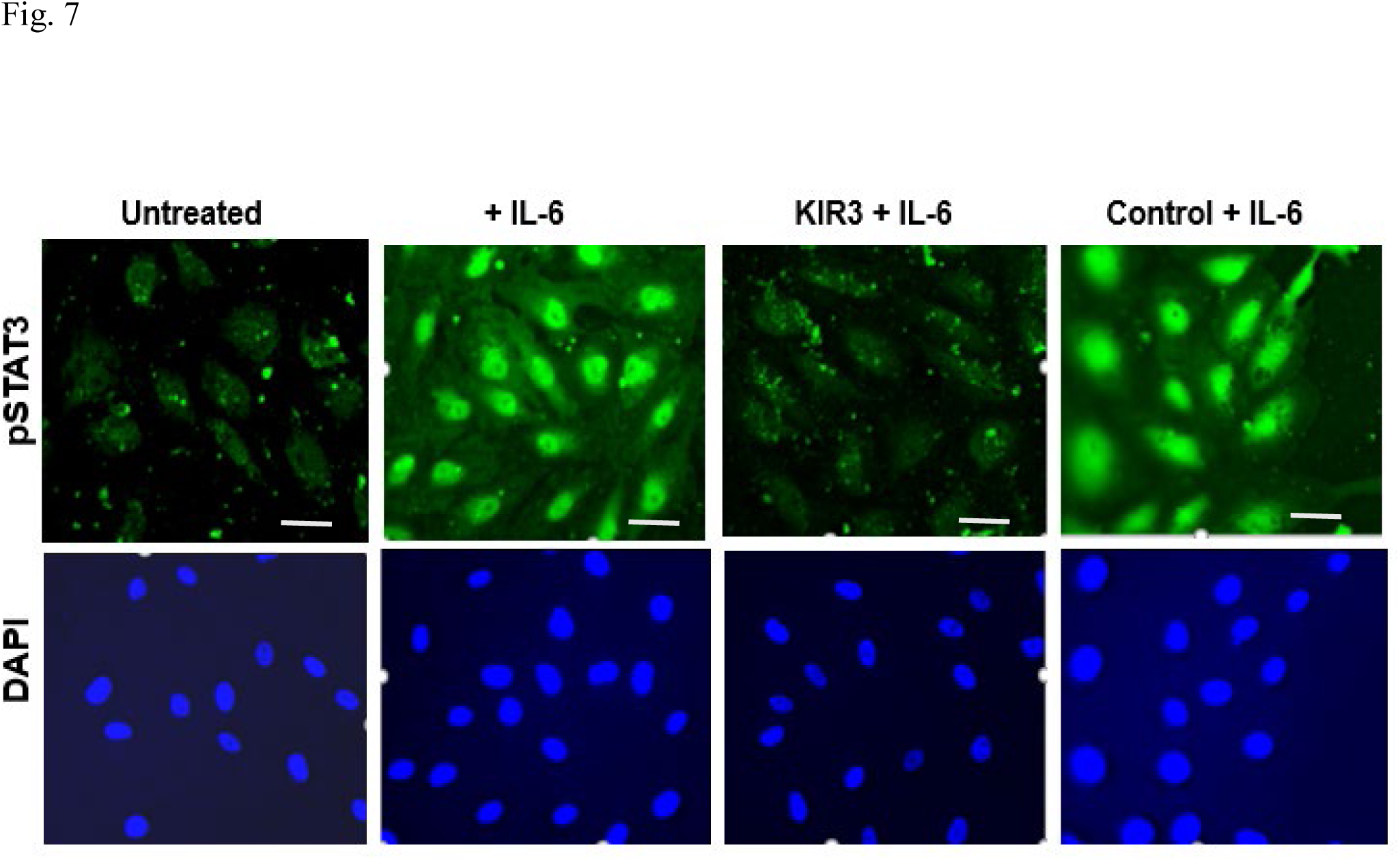
R9-SOCS3-KIR blocked IL-6 mediated nuclear translocation of pSTAT3. ARPE-19 cells in serum free medium were pre-treated R9-SOCS3-KIR (20 μM), or its control peptide for 1 hr followed by treatment with IL-6 (50 ng/ml) for 0.5 hr. Cells were washed and stained with antibody to pSTAT3, followed by staining with Alexa-488 conjugated secondary antibody and DAPI, and imaged with a fluorescence microscope. Scale bar, 50 μm.

### Corneal application of R9-SOCS3-KIR protects mouse eyes in the sodium iodate model

Having noted a combination of protective effects leading to alleviate the factors that result in AMD in the presence of R9-SOCS3-KIR in cell culture studies above, we tested its efficacy in protecting eyes from a sodium iodate (NaIO_3_) induced acute oxidative injury. Two groups of C57BL/6 mice (BomTac, n=10, both sexes) were pre-treated with eye drop administration of R9-SOCS3-KIR or its control peptide (15 μg in 2 μl) in both eyes, once a day on days –1 and 0. On day 0, mice were injected ip with 25 mg/kg of NaIO_3_. Daily eye drop instillation was continued for 3 days. On day 3, digital fundoscopy revealed influx of inflammatory cells, perivascular deposits and engorged blood vessels in the control group, while the R9-SOCS3-KIR treated mice had less damage (**Figure 8A**). Optical coherence tomography showed thinning of ONL and more of infiltrating cells in control treated eyes, as compared with the R9-SOCS3-KIR treated eyes (**Figure 8B**). On day 3, mice (n=4 in each group) were humanely sacrificed and their eyes were harvested, fixed and stained with hematoxylin and eosin (**Figure 8C**). A larger number of inflammatory cells in vitreous and retina were observed in controls, while the R9-SOCS3-KIR treated eyes had fewer of these cells. More notably, retinal swelling and rosettes (in-foldings) were a common feature of control eyes and were not found in any of the KIR3 treated eyes.

**Figure 8.**
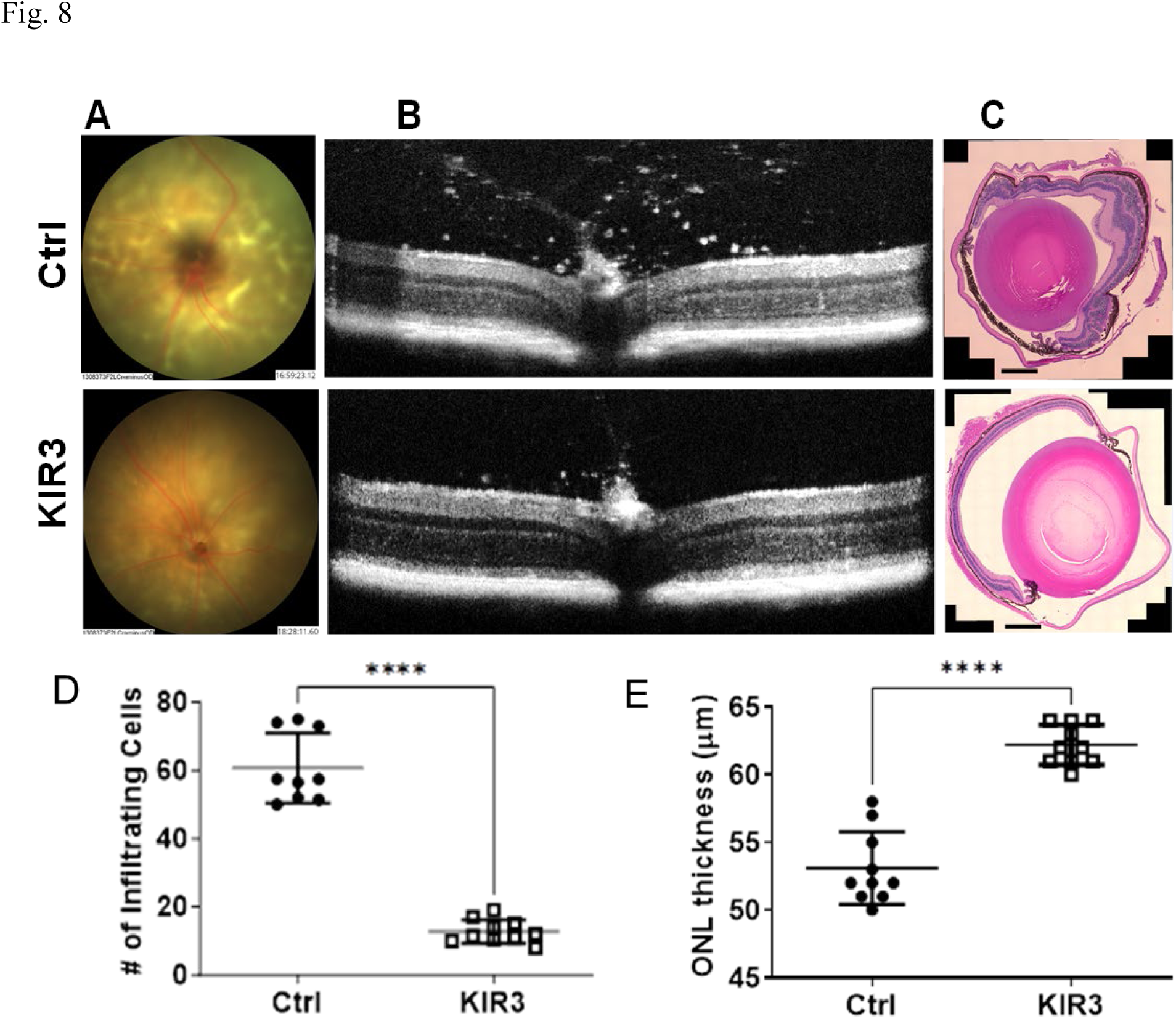
Corneal application of R9-SOCS3-KIR protected mouse eyes against injury caused by sodium iodate. C57BL/6 (BomTac, both male and female, n=10) were used for eye drop instillation of R9-SOCS3-KIR or its control peptide daily (15 μg in 2 μl in each eye) one day prior to injecting sodium iodate (25 mg/kg) intraperitoneally. Peptides were administered daily. On day 3, fundoscopy (**A**), and spectral domain optical coherence tomography (**B**) were carried out. H and E stained eyes from a control and a treatment eye (**C**); scale bar, 200 μm. The number of infiltrating cells (**D**) were counted using image J. The thickness of outer nuclear layer (ONL) was measured using the segmentation of Diver 2.0 of Leica microsystems (**E**). Units are micrometers. Statistical significance was tested using Student’s t test for unpaired data. ****, p<0.0001.

Using image J software, we quantified the number of inflammatory cells in B-scans from four areas of the posterior chamber from OCT images. In control eyes, we noted 60 ± 9 inflammatory cells per image, whereas the average number of cells in R9-SOCS3-KIR treated mice were 12.8 ± 3.4 (n=18, p < 0.001) per image, indicating less inflammation with R9-SOCS3-KIR (**Figure 8D**). The thickness of outer nuclear layer (ONL) was measured using the autosegmentation software Diver 2.0 of Leica microsystems (**Figure 8E**). The ONL in control eyes measured at 53.1 ± 2.6 (P = 0.001), while R9-SOS3-KIR treated eyes had ONL thickness of 62.2 ± 1.5 (p <0.001), indicating the protective role of R9-SOCS3-KIR. There was no significant difference between the male and female mice in the measurements noted above.

To investigate the protective effect of R9-SOCS3-KIR in mice eyes, we examined the induction of anti-oxidative genes. We harvested retina from control and treated mice (n=4 from each group) on day 3. RNA was extracted and used for reverse transcriptase followed by qPCR (**Figure 9**). We noted a 3-to-4-fold increase of heme oxygenase (HO-1), GSTM-1, NQO-1 and catalase in R9-SOCS3-KIR treated eyes, as compared with the control eyes. The antioxidant effect from these enzymes may have contributed to the protection in the eyes noted above.

**Figure 9.**
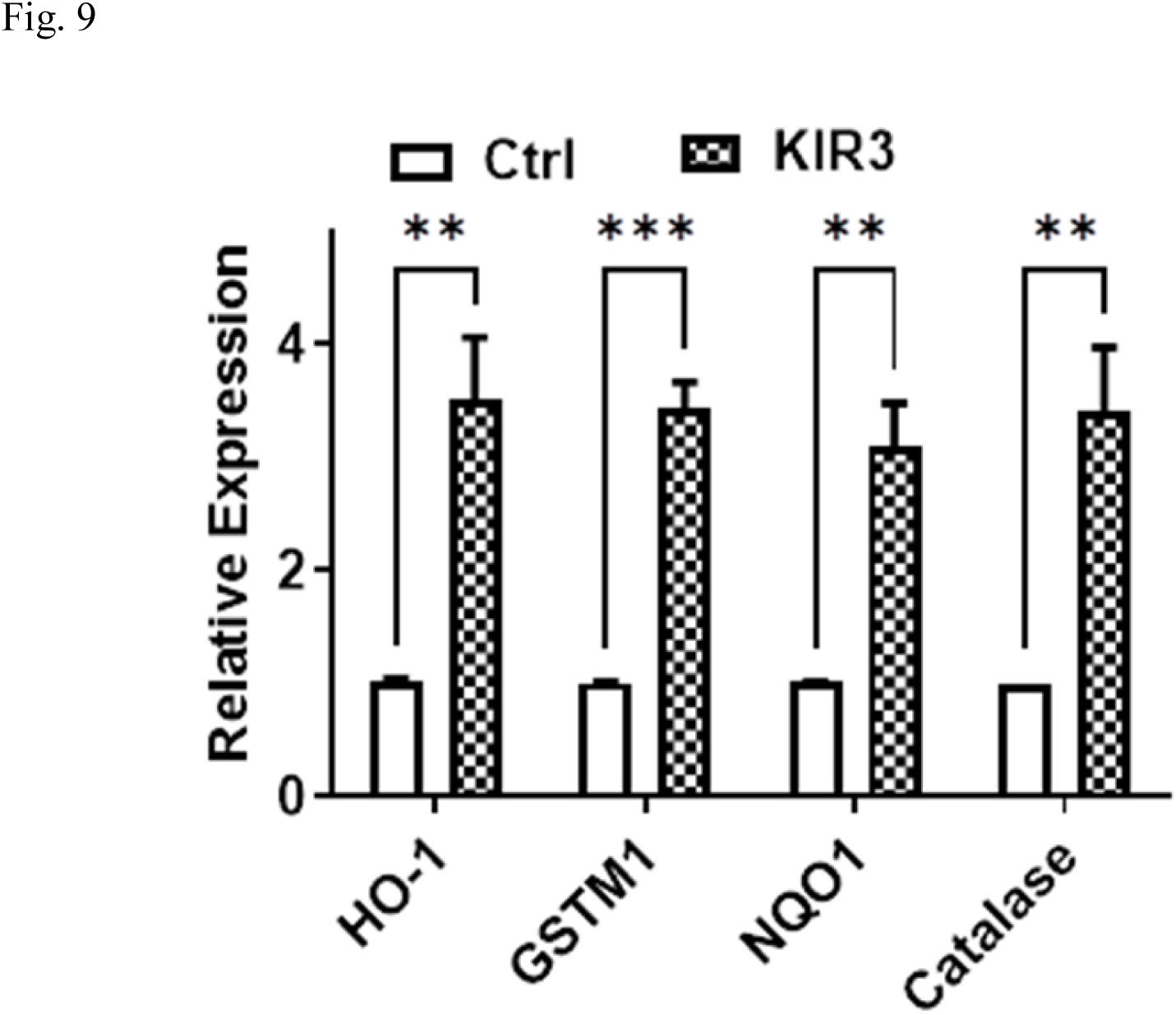
Eye drop instillation of SOCS3-KIR enhanced the anti-oxidant enzymes in retina of mice treated with sodium iodate. Mice (n=4 in each group) were treated on day –1 until day 3 with R9-SOCS3-KIR or control peptide (15 μg in 2 μl in each eye) and injected i.p. on day 0 with sodium iodate. On day 3, retina was isolated and used for RNA extraction and qRT-PCR with the target anti-oxidant enzymes and compared with β-actin as an internal control. In each comparison, Student’s t test for unpaired data was used to establish statistical significance for each comparison. **, p<0.01; ***, p<0.001. Abbreviations: HO-1, heme oxygenase 1; GSTM1, Glutathione S-Transferase Mu 1; NQO-1, NAD(P)H Quinone Dehydrogenase 1.

## DISCUSSION

R9-SOCS3-KIR is a water soluble, cell-penetrating peptide containing the kinase inhibitory region (KIR) of the human SOCS3 protein. Since this peptide is of human origin, there is no immunogenic response expected. It can be delivered as an eye drop, which limits its effect to the site of delivery and reduces the potential for systemic side effects, while allowing patient self-administration. We demonstrate its benefits against injurious processes that are interdependent and self-perpetuating: (1) dysregulated complement system that increases with age; (2) cytokine mediated inflammation that also intensifies with age; (3) elevation of VEGF-A that contributes to neovascularization; and (4) oxidative stress that may injure tissues directly and induce inflammation. The inhibition of these processes was accompanied by increased cell survival and junctional integrity of retinal pigment epithelial cells.

Inflammatory cytokines and growth factors that arise from an activated complement system were suppressed in the presence of R9-SOCS3-KIR. The NF-κB promoter that is central to the generation of the inflammatory mediators was suppressed as shown by the blockade of nuclear translocation of p65, an assay of promoter activity, retention of tight junction proteins, analysis of transcripts by qRT-PCR, and the suppression of VEGF-A release. The expression of cytokines IL-1β, IL-6 and IL-17A and the chemokine MCP-1 was suppressed in the presence of R9-SOCS3-KIR. IL-6 is known to compromise the immune privilege of the eye (48); thus its inhibition, should help maintain immune privilege. Oxidative stress has been shown to increase endogenous complement-mediated response, independent of the exogenously added complement source (49), suggesting these two processes to be interconnected and self-perpetuating. The membrane bound receptor for C5a and C3a is present on RPE and on several types of cells in the eye. The complement system is ordinarily a part of the innate arm of the immune system, but in AMD associated with CNV, splenic IL-17 producing γδT cells are transported to the eye (10, 50), thus the adaptive immune response also participates in the complement mediated visual impairment.

SOCS3-KIR peptide inhibits signaling downstream of the transcription factor STAT3 (**Figure 7**). This inhibition has implications in the signaling downstream from this transcription factor. Over-activation of STAT3 was shown to lead to elevated VEGF resulting in CNV (51). The optimal activation of STAT3 signaling is also important since persistent activation of STAT3 in retina induces visual impairment and retinal degeneration in aging mice (52). The SOCS3/STAT3 axis has been demonstrated by several investigators to be crucial in preventing various diseases (18, 23, 24, 53). In the same vein, Multiple Sclerosis (MS) patients on simvastatin therapy showed elevation in SOCS3 levels that resulted in decreased STAT3 phosphorylation and decreased IL-6 and IL-17 levels (54, 55). SOCS3 has been shown to prevent choroidal neovascularization in several studies (56–59) and has been shown to be a regulator of infection and inflammation (60, 61). Recently, the use of KIR peptide from SOCS3, attached to TAT protein domain for the purpose of cell penetration was reported in two publications from the same group for the suppression of IL-6 signaling in a model of traumatic brain injury (62, 63). The difference between the peptide we are presenting here and the one in these reports (62, 63) is that their KIR region started with the amino acid residue 22 and ended at 33, whereas the KIR3 region we are reporting extends from residues 20-35, and we have used 9 arginines for cell penetration.

SOCS1-KIR (26, 27) and SOCS3-KIR (described above) are specific and endogenous inhibitors of signaling from tyrosine kinases (TK, e.g. JAKs and TYKs), toll like receptor (TLR), and mitogen activated protein (MAP) kinases. The signaling from all of the preceding is mediated by the transcription factors, STAT1, STAT3, or NF-κB. A variety of tyrosine kinase inhibitors have been developed that are either monoclonal antibodies (mostly to the tyrosine kinases involved), or molecules screened from large-scale libraries with inhibitor properties. A recent example is the use of a monoclonal antibody to treat different mutations within the Myeloid-epithelial-reproductive (MER) tyrosine kinase (MERTK) that cause retinitis pigmentosa (RP) to varying degrees of severity (from mild to severe) or early in life loss of eyesight (64). A JAK1/JAK2 inhibitor ruxolitinib was therapeutic in this case. In case of traumatic brain injury (TBI), where MER tyrosine kinase was responsible in causing M1/M2 macrophage imbalance (65), TAM (Tyro3, Axl, and MER) receptor engagement was shown to absorb SOCS1 and SOCS3 (66). Thus, in cases in which MER tyrosine kinase is induced, R9-SOCS3-KIR might be a suitable treatment strategy.

Deficiency of SOCS3 in myeloid cells of mice led to prolonged activation of the JAK/STAT pathway and elevated expression of inflammatory cytokines such as IL-1β, IL-6, IL-12 and IL-23 (67). Conditional deletion of SOCS3 in blood and endothelial cells results in neutrophilia, pleural and peritoneal inflammation, and hematopoietic infiltration of several organs (68). In the retina, SOCS3 is expressed in photoreceptor cells. Conditional deletion of SOCS3 in photoreceptors suggests that it preserves the synthesis of rhodopsin, which is impaired following acute inflammation (69, 70). RPE cells also produce SOCS3 and SOCS1, and their levels are elevated in reaction to IL-17 and IFN-γ treatment, a response that would dampen an inflammatory response in the eye (71).

In the present experiments, we demonstrate that the kinase inhibitory region (KIR) of SOCS3 linked to a cell penetration sequence suppresses inflammatory signaling in two relevant cell types, RPE and macrophage. We also show that this peptide protects the integrity of the RPE monolayer following inflammatory insults (C5a and TNFα treatment) and protects RPE cells from mitochondrial oxidative stress induced by paraquat. Topical application of this peptide to the mouse cornea prevented injury to the retina caused by systemic delivery of sodium iodate in a model of acute retinal/RPE oxidative stress.

We have recently reported the corneal application of R9-SOCS1-KIR to treat endotoxin induced uveitis (27), and for the treatment of autoimmune uveitis as both a prophylactic and a therapeutic (26). Others have developed peptide therapeutics delivered topically to limit ocular angiogenesis and to provide neuroprotection (72–74). While there is considerable additional work required to establish an appropriate dose for human eyes and to determine the pharmacokinetics of R9-SOCS3-KIR and similar peptides, it seems likely that the topical use of peptide therapeutics can be developed for clinical practice.

## Acknowledgements

This work was supported by Shaler Richardson Professorship to ASL and by an unrestricted grant to the Department of Ophthalmology from Research to Prevent Blindness.

## Figure legends

**Supplemental Figure 1.**
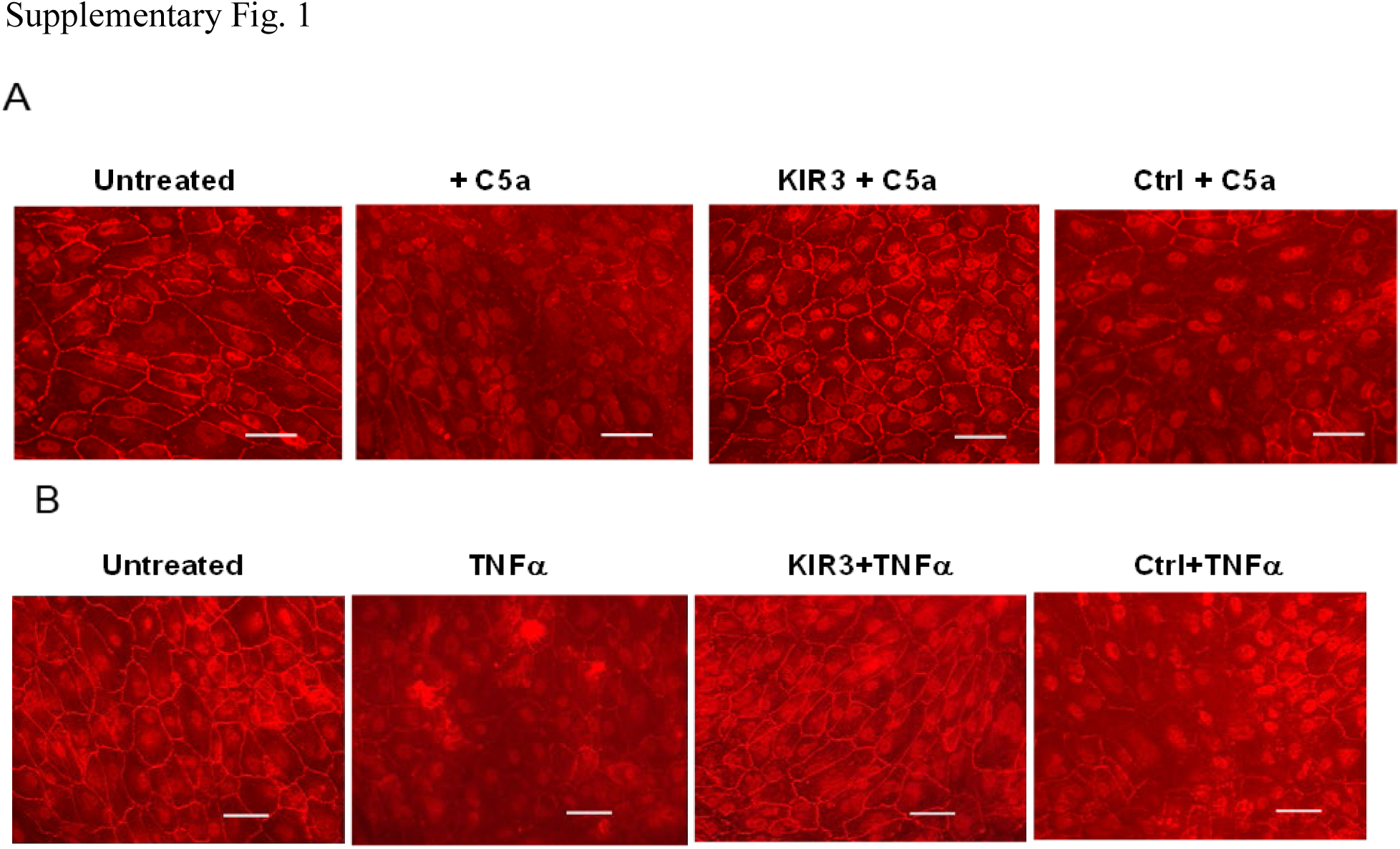
R9-SOCS3-KIR prevented loss of tight junction proteins caused by C5a or TNFα. ARPE-19 cells were grown in low serum media for 4 weeks until they had differentiated into a cobblestone monolayer. They were pretreated with R9-SOCS3-KIR or its control peptide (both at 20 μM) for 3 hr followed by treatment with **(A)** C5a (50 ng/ml) or (**B)** TNFα (10 ng/ml) for 48 hrs. Cells were permeabilized and treated with antibody to ZO-1 followed by incubation with a Cy-3 conjugated secondary antibody, washed, fixed and imaged using a Keyence fluorescence microscope. Scale bar is 50 μm.

**Supplemental Figure 2.**
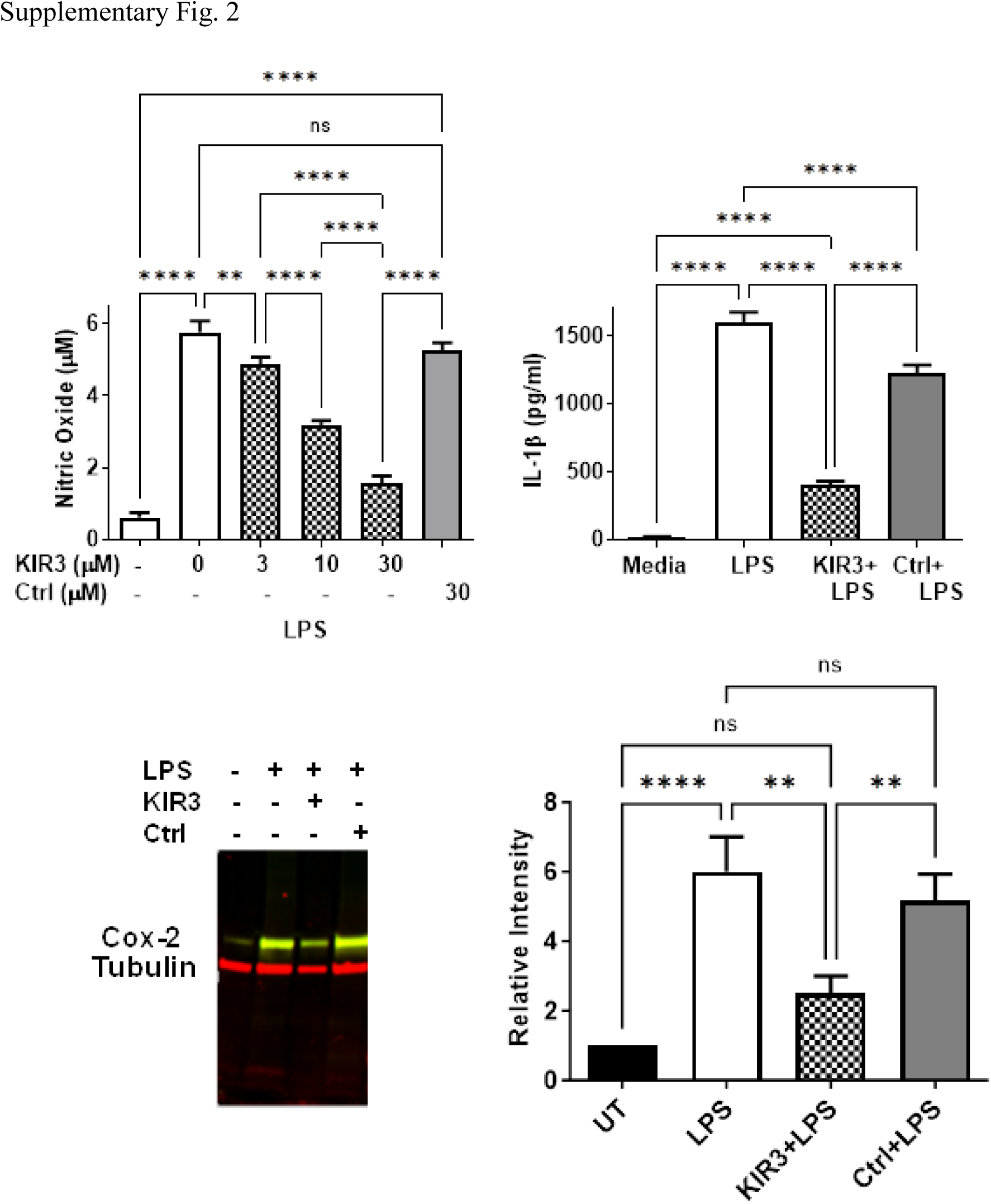
R9-SOCS3-KIR suppresses LPS mediated induction of NO production (A); decreases induction of IL-1β (B) and cyclooxygenase 2 (C and D). (**A**) Mouse macrophage cell line, J774.1 were pre-treated with indicated concentrations of R9-SOCS3-KIR or the control peptide (20 μM) for 1 hr, followed by addition of LPS (1 μg/ml) and the treatment overnight. Supernatants were harvested and used for NO estimation using Griess reagent. (**B**). J774A.1 cells were pre-treated with R9-SOCS3-KIR or its control peptide (20 μM) for 1 hr, followed by addition of LPS (1 μg/ml) and treatment overnight. Supernatants were harvested and used for quantitation of IL-1β by ELISA. One-way analysis of variance (ANOVA) followed by Tukey’s test for multiple comparisons showed significant differences. **, p < 0.01; ***, p<0.001; ****, p<0.0001. (**C**) J774A.1 treated as described were lysed to obtain cell extracts. Equal amounts of proteins were separated on a polyacrylamide gel, transferred to PVDF membrane and probed with antibodies to COX-2 and α-tubulin as an internal control. (**D**) The experiment as above was repeated three times for quantitation. The relative intensities of COX-2 and α-tubulin from three different blots were measured using image J software and averaged. One-way ANOVA followed by Tukey’s test for multiple comparisons showed significant differences between the treatments. **, p<0.01; ****, p<0.0001.

**Supplemental Figure 3.**
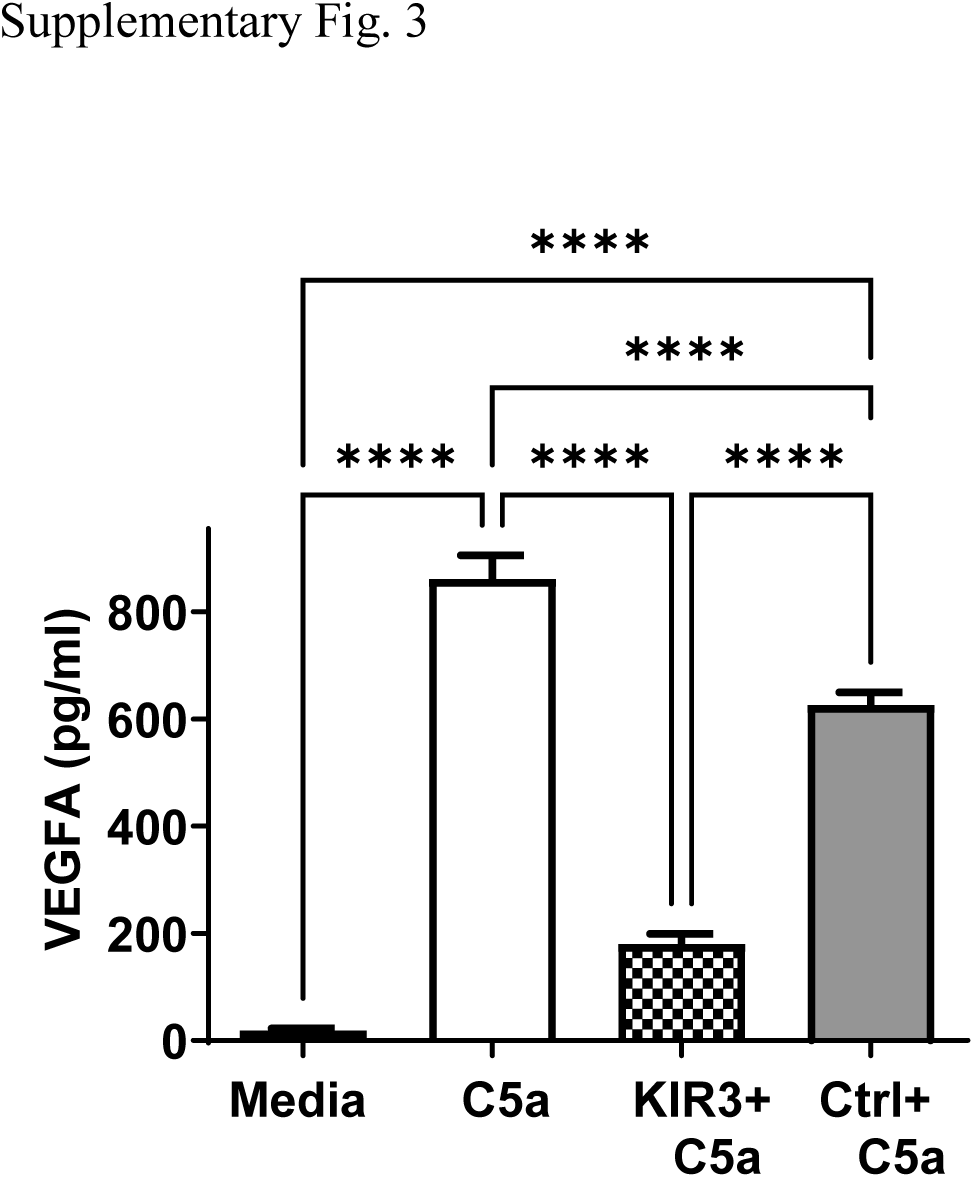
R9-SOCS3-KIR downregulated C5a-induced secretion of VEGF-A in J774A.1 cells. J774A.1 cells in 1% FBS containing media were treated with R9-SOCS3-KIR or the control peptide (at 20 μM) for 1 hr followed by treatment with C5a (50 ng/ml) and incubated overnight. Supernatants were harvested and used for quantitation of VEGF-A using a kit from PeproTech. ****, p <0.0001, as determined by one-way ANOVA.

